# Benchmarking of numerical integration methods for ODE models of biological systems

**DOI:** 10.1101/2020.09.03.268276

**Authors:** Philipp Städter, Yannik Schälte, Leonard Schmiester, Jan Hasenauer, Paul L. Stapor

**Affiliations:** Institute of Computational Biology, Helmholtz Zentrum München – German Research Center for Environmental Health, 85764 Neuherberg, Germany; Center for Mathematics, Technische Universität München, 85748 Garching, Germany; Faculty of Mathematics and Natural Sciences, University of Bonn, 53113 Bonn, Germany

## Abstract

Ordinary differential equation (ODE) models are a key tool to understand complex mechanisms in systems biology. These models are studied using various approaches, including stability and bifurcation analysis, but most frequently by numerical simulations. The number of required simulations is often large, e.g., when unknown parameters need to be inferred. This renders efficient and reliable numerical integration methods essential. However, these methods depend on various hyperparameters, which strongly impact the ODE solution. Despite this, and although hundreds of published ODE models are freely available in public databases, a thorough study that quantifies the impact of hyperparameters on the ODE solver in terms of accuracy and computation time is still missing. In this manuscript, we investigate which choices of algorithms and hyperparameters are generally favorable when dealing with ODE models arising from biological processes. To ensure a representative evaluation, we considered 167 published models. Our study provides evidence that most ODEs in computational biology are stiff, and we give guidelines for the choice of algorithms and hyperparameters. We anticipate that our results will help researchers in systems biology to choose appropriate numerical methods when dealing with ODE models.

## Introduction

Systems biology aims at understanding and predicting the behavior of complex biological processes through mathematical models [Kitano, 2002]. In particular, ordinary differential equations (ODEs) are widely used to gain a holistic understanding of the behaviour of such systems [Klipp et al., 2005]. These ODE models are often derived from biochemical reaction networks and stored/exchanged using the Systems Biology Markup Language (SBML) [Hucka et al., 2003] or Cell Markup Language (CellML) [Cuellar et al., 2003]. Based on these community standards, large collections of published ODE models have been made available to enhance reproducibility of scientific results, like BioModels [Li et al., 2010] and JWS [Olivier and Snoep, 2004]. These model collections allow an analysis of typical properties of published ODE models of biological processes and are an excellent source for method studies and method development [Hass et al., 2019].

Most published models of biochemical reaction networks are non-linear and closed form solutions are not available. Accordingly, numerical integration methods have to be employed to study them [Maiwald and Timmer, 2008]. For this task, the modeler has the choice between many possible numerical simulation algorithms, which have specific hyperparameters that can strongly impact the result. Key parameters are for instance the relative and absolute tolerance, which determine the precision of the numerical solution. Too relaxed tolerances may lead to incorrect results, while too strict tolerances may result in an unnecessarily high computation time, or may even lead to a failure of ODE integration, if the desired solution accuracy cannot be achieved [Hindmarsh et al., 2005].

Various theoretical results about the reliability and the scaling behaviour of ODE solvers are available [Ascher and Petzold, 1998]. However, to the best of our knowledge, there is no comprehensive study on the impact of ODE solver settings on the simulation results and their reliability which focuses on models of biological processes. So far, case studies using only single models or a very small number of models have been carried out, which demonstrate the need for efficient implementations of ODE solvers (see, e.g., [Maiwald and Timmer, 2008]). In addition to this, various hypotheses on the general properties of ODE models in systems biology exist, e.g., whether or not the underling ODEs are expected to be stiff – an ODE is (informally) called stiff, if it exhibits different time-scales, i.e., fast and slow dynamics are described at the same time [Mendes et al., 2009, Raue et al., 2013]. The absence of representative studies and statistical evaluations is surprising as various tasks require large number of numerical simulations, rendering computation efficiency and numerical robustness an important topic. In particular when performing parameter estimation, a model has to be simulated, i.e., the underlying ODE has to be solved, thousands to millions of times [Maiwald and Timmer, 2008, Raue et al., 2013]. Indeed, it has recently been pointed out that the ODE solver is actually a crucial hyperparameter, which often remains unconsidered [Kreutz, 2019]. Hence, studying these questions on a wide set of real world applications is of high importance for many modeling applications.

In this work, we benchmark numerical integration methods and their hyperparameters. We established a benchmark collection of 167 models from the two freely accessible databases BioModels [Li et al., 2010, Malik-Sheriff et al., 2019] and JWS Online [Olivier and Snoep, 2004], which covers a broad range of different properties. These models were simulated using various ODE solver algorithms implemented in the SUNDIALS package CVODES [Cohen et al., 1996, Hindmarsh et al., 2005], a widely used state-of-the-art toolbox for solving differential equations, which offers a set of different methods. We investigated various combinations of ODE integration algorithm, non-linear solver employed in implicit multi-step methods, linear solvers employed within the non-linear solver, and relative and absolute error tolerances. By analyzing the computation time and the failure rate, we derived guidelines for the tuning of ODE solvers in systems biology, which facilitate fast and reliable simulation of the corresponding ODE systems.

## Results

To analyze combinations of algorithms and hyperparameters, we considered the ODE solvers implemented in SUNDIALS package CVODES [Cohen et al., 1996, Hindmarsh et al., 2005]. CVODES is used in multiple systems biology toolboxes [Fröhlich et al., 2017a, Raue et al., 2015] and is therefore particularly relevant. It implements implicit multi-step methods for numerically solving an initial value problem, i.e., an ODE with initial conditions. These algorithms offer a variety of hyperparameters. An initial value problem is solved by iterative time-stepping, following a specific integration algorithm [Ascher and Petzold, 1998], (see Methods, Numerical integration methods for ODEs for more details). In each time step, a non-linear problem is solved via a fixed-point iteration or a sequence of linear problems. These are solved until a previously defined precision, given by absolute and relative error tolerances, is fulfilled.

CVODES offers the following hyperparameters:

1. Integration algorithm:
  a. Adams-Moulton (AM): implicit multi-step method of order 1 to 12
  b. Backward Differentiation Formula (BDF): implicit multi-step method of order 1 to 5
2. Non-linear solvers:
  a. Functional: solution to the non-linear problem directly via a fixed-point method
  b. Newton-type: linearization of the non-linear problem
3. Linear solver (only when using Newton-type non-linear solver):
  a. DENSE: dense LU decomposition
  b. GMRES: iterative generalized minimal residual method on Krylov subspaces
  c. BiCGstab: iterative biconjugate gradient method on Krylov subspaces
  d. TFQMR: iterative quasi-minimal residual method on Krylov subspaces
  e. KLU: sparse LU decomposition
4. Error tolerances: upper bounds for the absolute and relative error made in each time-step

We call a combination of these hyperparameters a solver setting.

As it is still unclear which solver settings are best suited for models of biochemical reaction networks, we performed an exhaustive empirical study to answer this question. Therefore, we considered all 20 possible combinations of integration algorithm, non-linear solver, and linear solver. Furthermore, we considered 36 error tolerance combinations in an in-depth tolerance study, and 7 out of those 36 in the remaining studies. This yielded in total 169 different solver settings.

As performance characteristics of a solver setting, we consider the following two criteria:

1. Integration failures: These failures may occur if either the dynamics of the system become too stiff or if the state of the system diverges. In these cases, the requested numerical accuracy per integration step cannot be achieved by a solver setting and the (adaptively chosen) step-size falls below machine precision and integration gets stuck.
2. Computation times: The total computation time is determined by the number of steps the solver takes and their individual computation times. Both quantities can vary heavily depending on the solver settings.

### A comprehensive model collection allows the systematic bench-marking of hyperparameters

For a comprehensive study on a variety of ODE models, we downloaded all SBML models from the JWS database [Olivier and Snoep, 2004]. As all of those models had less than 100 state variables, we complemented them with a set of the largest models from the Biomodels database [Li et al., 2010]. To simulate these models using CVODES, we imported them with AMICI (Advanced Multi-language Interface to CVODES and IDAS), which performs symbolic preprocessing and creates and compiles executable code for each model. As AMICI does not support all features of SBML (such as assignment rules for state variables or discrete events), the import worked only for a subset of these models. To ensure a proper comparison, we verified the correctness of the simulated trajectories (Figure 1A), based on reference solutions. After this filtering step, we were left with 167 ODE models, of which a majority comprises between 10 and 100 state variables and reactions (Figure 1B). For details on the construction of the benchmarking collection (e.g., source of reference solutions) we refer to the Methods, Creation of the ODE solver benchmark collection.

**Figure 1:**
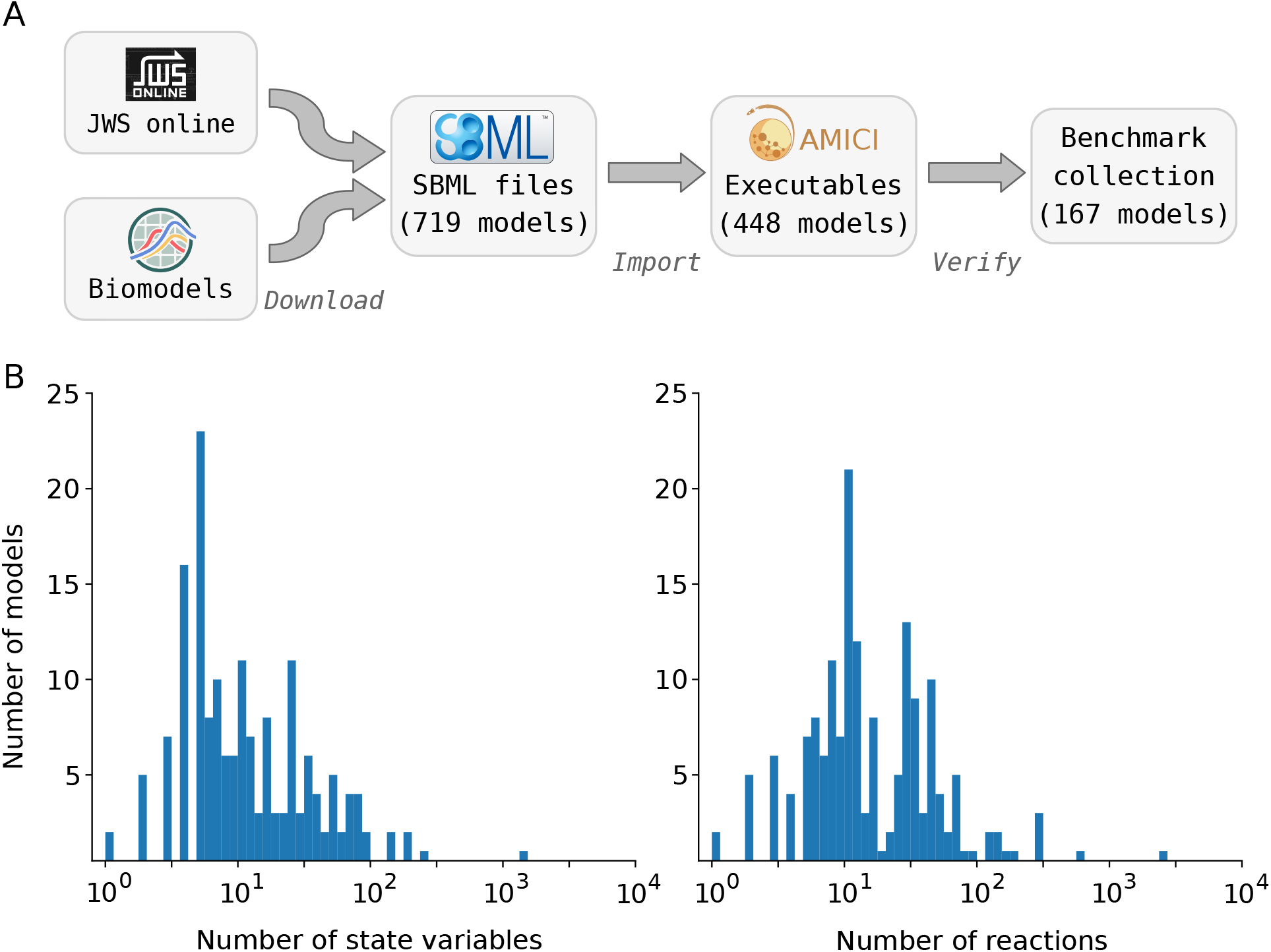
Model Collection. **A:** Workflow of collecting models for the benchmark collection. Models were downloaded, imported with AMICI, and simulation results were compared to reference trajectories using multiple solver settings. **B:** Basic properties (number of state variables and reactions) of the benchmark collection models.

### Newton-type method outperforms functional iterator when solving the non-linear problem

Using the 167 ODE models, we analyzed the impact of different hyperparameters on the performance of numerical ODE solvers. As we expected the combination of the non-linear and linear solver to strongly impact the reliability of the numerical ODE integration, we decided to first study this aspect.

We compared the rate of integration failure for the two available non-linear solvers – the functional iterator and the Newton-type solver – for both ODE integration algorithms (AM and BDF), all available linear solvers and a set of heterogeneous error tolerances, motivated by commonly used ODE solver settings. This comparison revealed that the Newton-type method performs better than the functional iterator: While the functional iterator, which does not rely on a linear solver, failed on average for 10 – 15% of the models, the Newton-type method failed substantially less (Figure 2). Indeed, in combination with the BDF integration algorithm, we observed for all but the dense direct linear solver DENSE a failure rate of less than 5% of the models. Overall, the BDF method appeared to be less prone to integration failure than the AM method. This result can be considered as hint that the models are stiff, as the AM method is less tailored to models exhibiting stiff dynamics [Hairer and Wanner, 1996]. Interestingly, the highest failure rate was observed for the Newton-type method in combination with the dense linear solver. For all other linear solvers, the Newton-type method is less prone to integration failure than the functional iterator and also more efficient in terms of computation time (Supplementary Figure S2). Thus, we focus in the following only on results for the Newton-type method.

**Figure 2:**
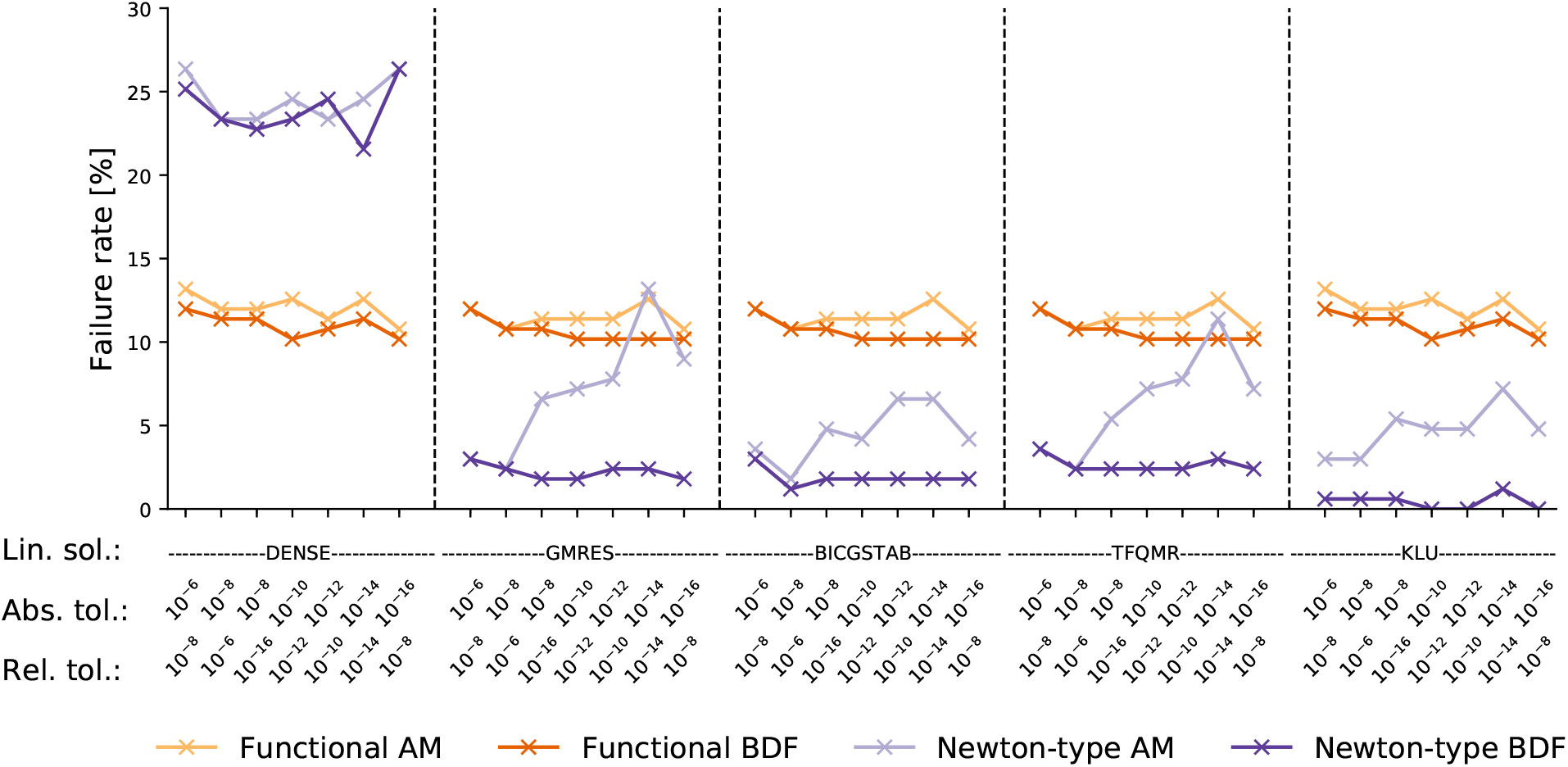
Non-linear solver. Comparison of Functional and Newton-type non-linear solver in terms of failure rate. Each point represents the failure rate when simulating all models for a given combination of integration algorithm, non-linear solver, linear solver and error tolerances.

### The sparse direct linear solver scales best

To determine the performance of the different linear solvers, we assessed their computation times. For the BDF algorithm, we found that the dense direct solver exhibited the worst scaling behavior with respect to the number of state variables, followed by the iterative solver TFQMR (Figure 3A). Those two linear solvers also failed to integrate the largest models of the benchmark collection (Supplementary Figure S4), i.e., a cancer signaling model with 1228 state variables. The iterative linear solvers GMRES and BICGSTAB showed a sublinear scaling behavior for the BDF algorithm, i.e., when increasing the model size five-fold, the computation time increased on average roughly four-fold. The sparse direct solver KLU had the best scaling behavior. For the BDF method with the linear solver KLU, the complexity of the numerical integration seemed to grow approximately with the square root of the number of state variables.

**Figure 3:**
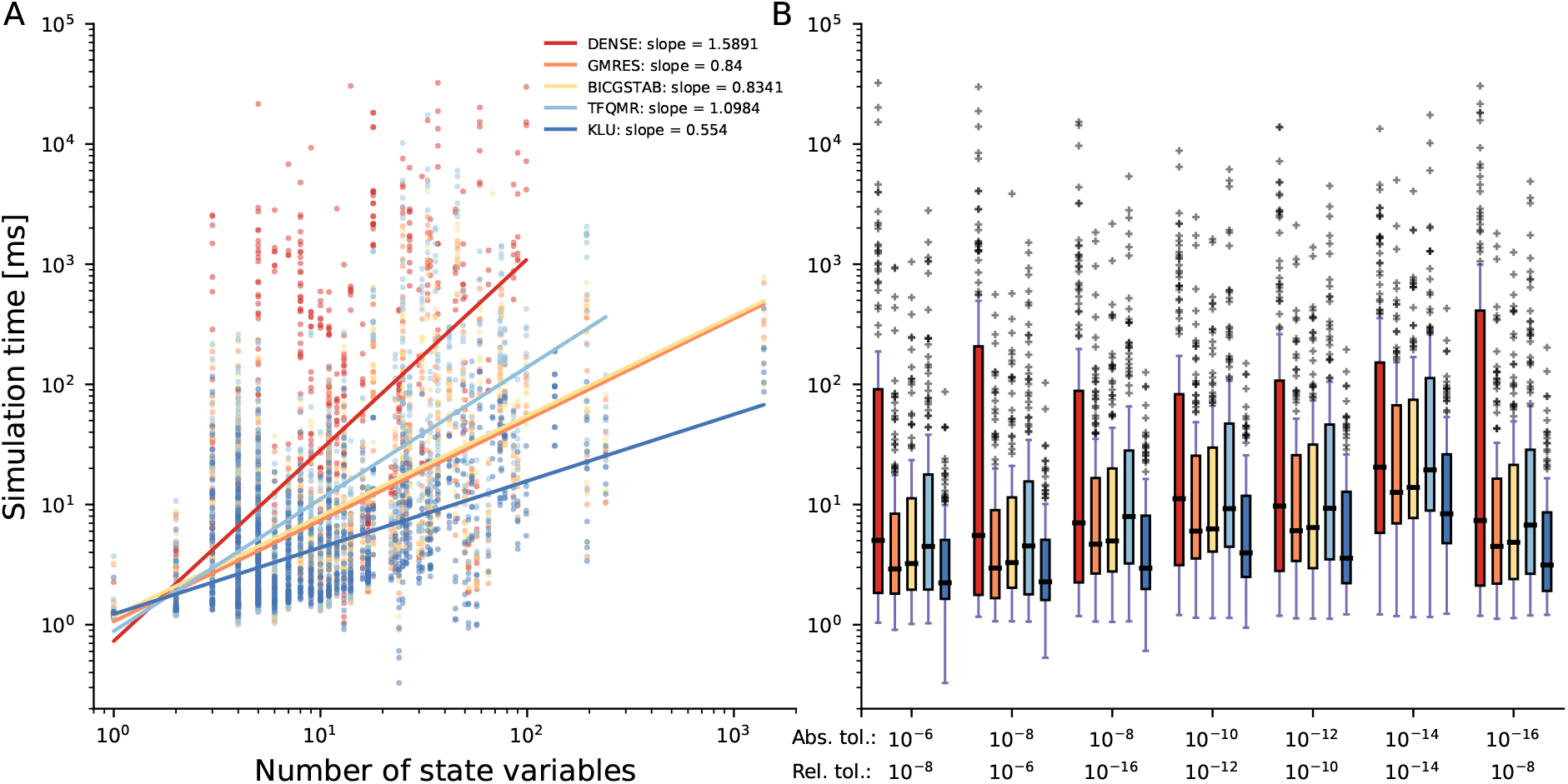
Linear Solvers. Computation time comparison of the five linear solvers. **A:** Each point depicts the simulation time of one model (median of 40 repetitions) with one solver setting: BDF integration algorithm, Newton-type non-linear solver, one tolerance combination, and one linear solver (which are color-coded). Results are shown for seven tolerance combinations and five linear solvers, meaning there are 35 points for each model. The accompanying linear regressions display the scaling behavior with respect to the number of state variables. **B:** Box plot of the simulation times, separated by the tolerance combination in addition to the linear solver.

Overall, the scaling behavior of most methods was slightly worse for the AM than for the BDF integration algorithm, but the overall results were similar: The computational complexity for the linear solvers GMRES and BICGSTAB increased linearly with the number of state variables, and the linear solvers TFQMR and DENSE showed the worst and KLU the best performance (Supplementary Figure S3).

Assessing the computation times across models for different error tolerances confirmed that the sparse direct linear solver KLU performed best for every tolerance combination (Figure 3B, Supplementary Figure S4). In contrast, the dense linear solver DENSE yielded the highest computation times. Concerning the three iterative linear solvers, minor differences in the computation time for GMRES and BICGSTAB were found, as GMRES was slightly faster than BICGSTAB. As expected, stricter error tolerances consistently led to an increase of computation time for all employed solver settings.

### Choosing error tolerances is a trade-off between accuracy, reliability, and computation time

In a next step, we studied the effect of the error tolerances on the computation time. As the Newton-type non-linear solver with the sparse direct linear solver KLU outperformed all other combinations, we used this combination to analyze the impact of the absolute and relative error tolerances on computation time and failure rate. For an extensive tolerance study, not only the hitherto seven, but 36 tolerance combinations were analyzed, covering a broad spectrum. As upper bounds for the relative and absolute tolerances we used 10^−6^, as more relaxed tolerances provided an insufficient agreement with the reference trajectories (see Methods, Creation of the ODE solver benchmark collection and Supplementary Figure S1).

To compare computation times across all models, we analyzed CPU time ratios. The computation time for each model and each error tolerance combination was normalized by the computation time for the most relaxed tolerance combination, i.e., absolute and relative tolerance of 10^−6^ (Figure 4A, Supplementary Figure S5). We found that for most models, the computation time increases with the enforced accuracy. Yet, we found a few outliers, indicating a non-trivial relation. Apart from those exceptions, the CPU time indeed increased monotonically when restricting the absolute and relative error tolerances to 10^−16^: The median computation time increased by slightly more than a 6-fold. Hence, restricting the requested error tolerances by ten orders of magnitude increased the median computation time by less than one order of magnitude, which was less than we had expected. Furthermore, strict relative error tolerances tended to have a bigger impact on the computation time than strict absolute tolerances.

**Figure 4:**
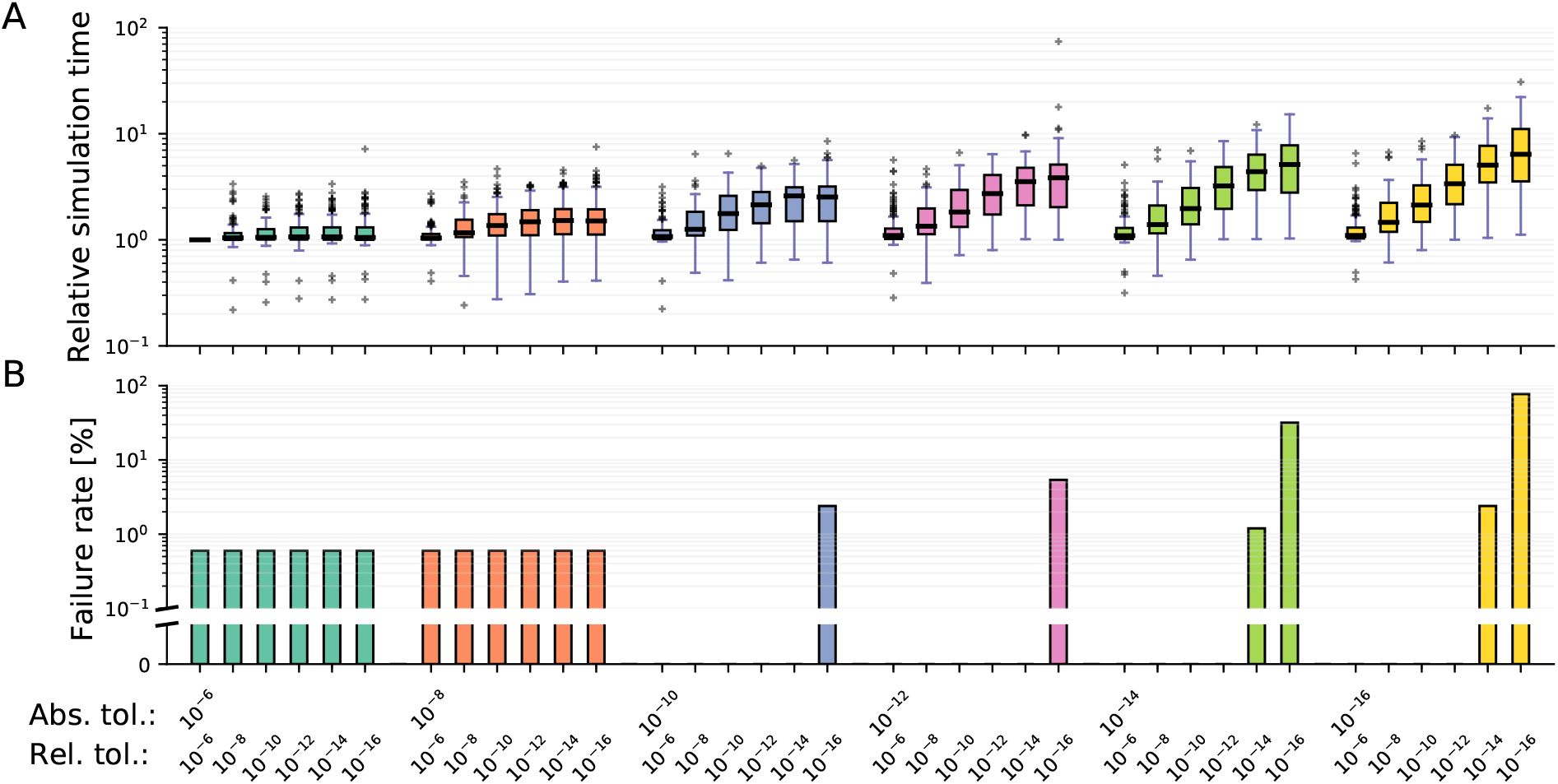
Integration error tolerances. Comparison of 36 combinations of absolute and relative error tolerances. All models were simulated using the BDF integration algorithm, the Newton-type non-linear solver, and the linear solver KLU, for each tolerance combination. **A:** Relative simulation times, normalised by those for absolute and relative error tolerances (10^−6^, 10^−6^). **B:** The corresponding failure rates.

The analysis of the failure rate with respect to the error tolerances revealed that relative error tolerances impacted integration problems to a higher degree than absolute error tolerances (Figure 4B): E.g., when comparing the tolerance combination for the absolute and relative tolerances (10^−16^, 10^−12^) with the inverse combination, the former allowed to simulate all models, whereas the latter failed for 7 models. However, substantial increases in the failure rate only occurred when both error tolerances were restricted at the same time.

### Fully implicit algorithms are the best choice for most models

Since we have seen noticeable differences in their behavior, we compared the integration algorithms AM and BDF for each model individually regarding their computation time and failure rate. Therefore, we used again the seven tolerance combinations which we employed for the analysis of the non-linear and the linear integration algorithms.

We observed that for most models and settings, the BDF algorithm was faster than the AM algorithm (60% vs. 36%, Figure 5A). For a number of models, BDF was faster by almost two orders of magnitude when compared to AM. In contrast, AM outperformed BDF by at most a factor of 4 (Figure 5B). For small models with less than 10 state variables, AM tended to be faster, while BDF clearly outperformed AM for larger models.

**Figure 5:**
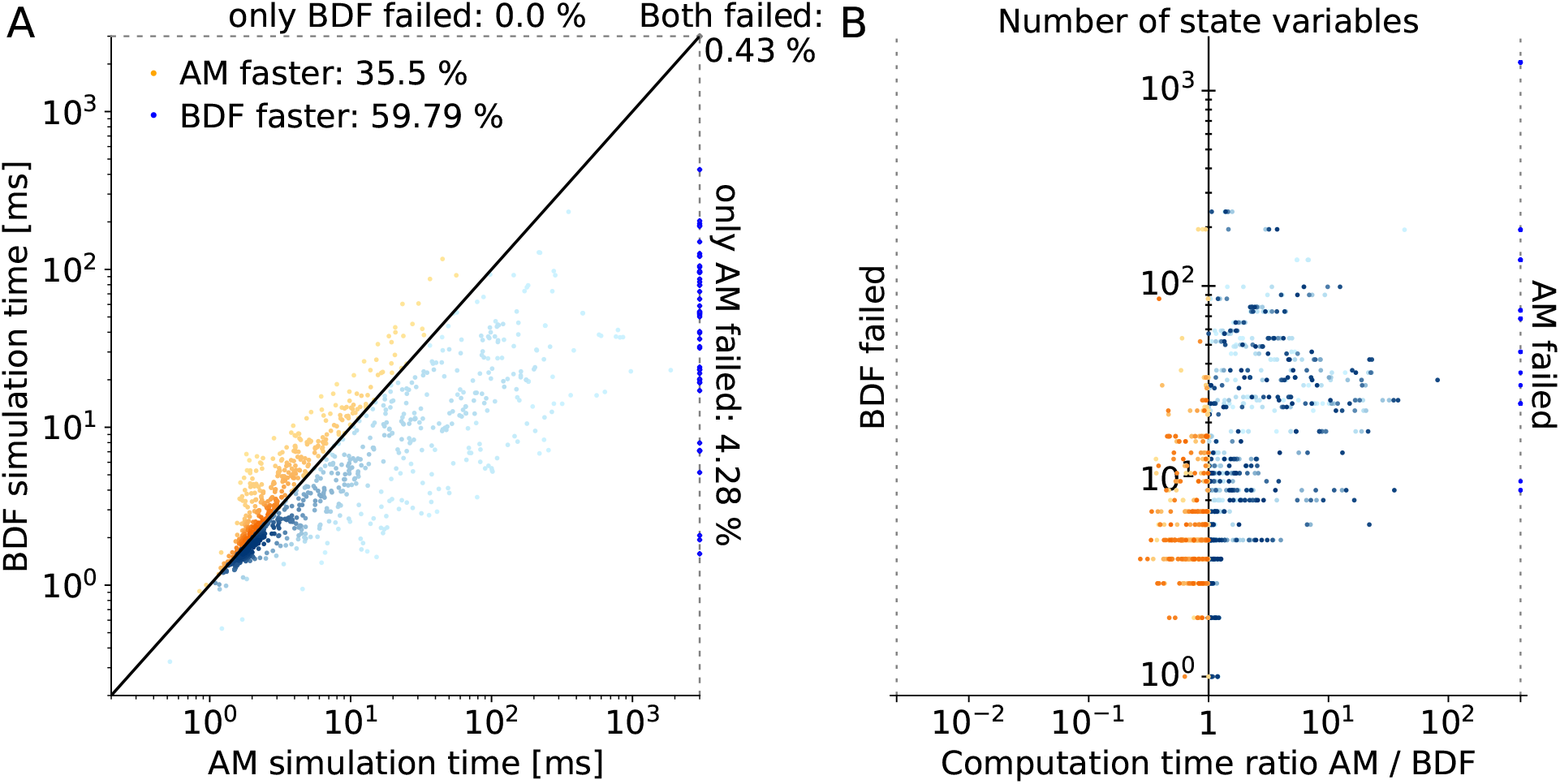
Integration algorithm AM vs. BDF. Comparison between AM and BDF integration algorithm in terms of failure rate and computation time. **A:** Each scatter point represents the computation time for the numerical integration of a model using AM (x-axis) or BDF (y-axis) in combination with the Newton-type non-linear solver, the linear solver KLU and one tolerance combination. The results for all seven original tolerances are shown, meaning that there are seven points for each models. The scatter points on dotted lines on top and on the right denote failed simulations with only one or both integration algorithms. Darker colors represent a higher scatter point density. **B:** Computation time for AM divided by the computation time for BDF with respect to the number of state variables, using the color coding from subfigure A.

Importantly, the BDF algorithm was not only computationally more efficient, but showed also the lower failure rate: For about 0.4% of the settings, both algorithms failed to integrate the ODE. For an additional 4% of the cases, BDF could integrate the ODE while AM failed. Hence, the AM algorithm failed about 10 times more often to integrate the ODE system than the BDF algorithm.

When comparing the two integration algorithms AM and BDF with the remaining four linear solvers, we found a similar situation (Supplementary Figure S6): BDF could integrate more models and settings than AM and was overall the faster method. Both algorithms failed more often when not using the sparse linear solver KLU. This again underlines that the linear solver KLU outperforms the other linear solvers in terms of reliability. Afterwards, we additionally included the functional non-linear solver (Supplementary Figure S7): Also in this case, BDF could integrate more models and settings than AM and was still overall the faster method, and the overall failure rate increased due to including the functional iterator.

## Discussion

Modeling with ODEs is among the most popular approaches to develop a holistic understanding of cellular processes in systems biology. Here, we collected a set of 167 published ODE models and used them to carry out a comprehensive study on the most essential hyperparameters of numerical ODE solvers. To the best of our knowledge, this is the first extensive study focusing on ODE integration itself, investigating a total of 169 ODE solver settings. The use of a large number of established models makes it highly relevant to the community. Although the optimal choice of hyperparameters is model dependent, our findings allow to draw some general conclusions:

Firstly, we found that, in general, error tolerances should not be relaxed beyond the value of 10^−6^, as otherwise simulation results tend to deviate markedly from results of more accurate computations. However, too strict error tolerances substantially increase the computation time and, more importantly, can lead to failure of ODE integration. We conclude that for most models, error tolerances between 10^−8^ and 10^−14^ are a reasonable choice, with absolute error tolerances being stricter than relative error tolerances, at least for the ODE solver implementation considered in this study.

Secondly, we observed that for more than 64% of the models, the BDF integration algorithm was superior to the AM integration algorithm. As the AM algorithm is generally recommended for mildly stiff problems and BDF is more tailored to stiff systems [Hairer and Wanner, 1996], this implies that most ODE models in systems biology show substantial stiffness. Stiffness was already hypothesized for ODEs arising from biological systems [Klipp et al., 2005, Mendes et al., 2009, Raue et al., 2013], but to the best of our knowledge, this was never quantified. Hence, our results suggest that fully implicit methods for ODE integration with adaptive time-stepping and error control are necessary to obtain reliable results, except for special cases, where clear motivations for other approaches can be given. We also want to stress that all ODE models were analyzed at the reported parameter values, for which ODE integration is supposed to work well. However, these parameters often have to be estimated first, by evaluating the ODE at various parameters and comparing model simulations with measurement data [Raue et al., 2013, Villaverde et al., 2018]. During this estimation process, the model does often not yet reflect a realistic behavior and in our experience stiffness is encountered substantially more often. Hence, in parameter estimation, stiffness is likely to be even more present.

Thirdly, when comparing algorithms for solving the non-linear and the linear problem within implicit integration methods, we found that the fastest and most reliable setting is using a Newton-type method for the non-linear problem and a sparse direct solver for the linear problem (in our case KLU), in particular with a better scaling behavior towards higher-dimensional models. This setting performed best, independent of the algorithm employed for integrating the ODE. In contrast, the dense direct linear solver showed the worst performance. Furthermore, it was pointed out that for stiff ODE systems, Newton-type non-linear solvers tend to be superior to fixed-point methods [Norsett and Thomsen, 1986], which we can underline by our findings.

While this study focused on numerically solving ODE systems, a valuable extension would be assessing the performance of ODE integration when performing it with forward and adjoint sensitivity analysis [Fröhlich et al., 2017a]. This is a typical setting when estimating unknown model parameters, as sensitivities are needed to compute the gradient of an objective function which depends on the ODE solution [Raue et al., 2013]. It would be particularly interesting to see whether it could be inferred when forward and when adjoint sensitivity analysis should be used: Although forward sensitivity analysis is more commonly used and may be superior for small models, adjoint sensitivity analysis is known to be more efficient for large models [Sengupta et al., 2014, Frohlich et al., 2017a]. Yet, no clear rule for deciding between those two approaches could be derived so far. The respective sensitivity equations may also change in particular the stiffness of the ODE, which may impact optimal hyperparameters.

Moreover, the multi-step algorithms in CVODES could be compared to other approaches, such as the (implicit) single-step algorithms which are provided in, e.g., the toolbox FATODE [Zhang and Sandu, 2014]. A comparison between CVODES and LSODA [Petzold, 1983] might also be interesting. LSODA automatically chooses between the AM and the BDF integration algorithms depending on the degree of stiffness for the current ODE. However, its probably most widely used implementation is not actively maintained anymore and does not support the simultaneous integration of sensitivity equations.

In conclusion, we are certain that the presented study will be helpful for both modelers and toolbox developers, when choosing hyperparameters for ODE solvers to yield both reliable and efficient simulations.

## Methods

### Numerical integration methods for ODEs

An ODE model describes the dynamics of a vector of state variables 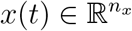 with respect to time *t*, model parameters *θ*, and input parameters *u*. The time evolution is given by a vector field *f*:

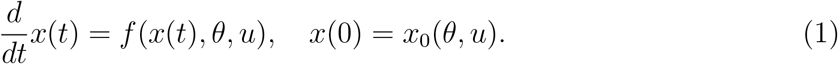

As typically the solution *x*(*t*) cannot be computed analytically, we consider in this study different numerical integration methods which proceed along the time course step by step: Starting from time *t*_0_ and state *x*_0_, an approximation *x_k_* of the state *x*(*t_k_*) at the *k*-th time-step *t_k_* is computed from the states and the vector fields at previous time points. In so-called multi-step methods, the *s* previous time-steps of the ODE solver *t*_*k*–1_, …, *t*_*k*–*s*_ are used, where 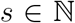 fixes the order of the method. The most general form of a multi-step method is:

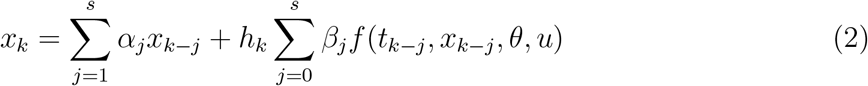

The difference *h_k_* = *t_k_* – *t*_*k*–1_ defines the time-step length, and the coefficients *α_j_* and *β_j_* determine the exact algorithm. When *β*_0_ = 0, the method is explicit, otherwise implicit. Generally, implicit algorithms are considered to be better suited for ODE systems with so-called stiff dynamics [Hairer and Wanner, 1996].

For implicit methods and non-linear vector fields *f*, a non-linear problem has to be solved to determine *x_k_*. In order to solve the non-linear problem, either a fixed-point iteration can be used, which tries to approximate *f*(*t_k_*, *x*(*t_k_*), *θ*, *u*) step by step, or a Newton-type method can be employed, which reduces Equation (2) to a series of linear problems, which can be denoted as:

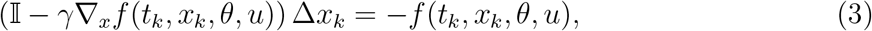

where is the unit matrix 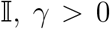 a relaxation constant and ∇*_x_f*(*t_k_*, *x_k_*, *θ*, *u*) the Jacobian of the right hand side. In the latter case, algorithms for solving linear problems have to be used repeatedly. Here, we consider two approaches: Direct approaches, which try to solve Equation (3) by factorization of the matrix *A* (such as LU factorization); and iterative approaches, such as Krylov subspace methods, which try to compute *x* by a sequence of improving approximations *x*^0^, *x*^1^,…, *x^m^*, until *x^m^* solves Equation (3) up to a previously defined accuracy.

### High performing software packages for ODE integration

It has been shown in several studies that it is critical to use high-performance ODE solver toolboxes written in compiled languages such as C++ or FORTRAN, since those have shown to reduce computation time by two to three orders of magnitude [Maiwald and Timmer, 2008, Kapfer et al., 2019]. For this reason, we focused on the ODE solvers in CVODES, which is part of the SUNDIALS solver package [Hindmarsh et al., 2005]. These C-based solvers are used in several established toolboxes, e.g. Data2Dynamics [Raue et al., 2015] or AMICI [Fröhlich et al., 2017b]. Appropriate alternatives might be in our opinion other solver toolboxes written in compiled languages, such as the FORTRAN-based FATODE implementation [Zhang and Sandu, 2014] or the FORTRAN-based ODEPACK implementation [Hindmarsh, 1983], or reimplementations of these.

### Setup of the numerical experiments

All numerical simulations were repeated 40 times to avoid outliers in the computation time. Afterwards, the median values of all these computation times were stored. Since a repeated simulation using the same solver setting does not change the success of the simulation, this information was stored after the first simulation. For the models from the JWS database, the time interval consisting of start time, end time and time steps for simulating a model was taken from the accompanying SED-ML file. For the remaining models, where no SED-ML files with such specifications were available, 50 time steps were used, while the end points were inferred by visual inspection of the trajectories and set to values such that all state trajectories had reached a steady state. In total, 23,380 data points for computation time and success of the simulation were created. These were then analyzed and displayed in different ways to illustrate the impact of the hyperparameters on simulation time and reliability. The whole dataset is made available at https://doi.org/10.5281/zenodo.4013853. The study was performed on a laptop with Ubuntu 16.04.6 LTS operating system, Intel(R) Core(TM) CPU with maximal capacity of 2.80GHz, and 8GB RAM.

### Creation of the ODE solver benchmark collection

As a first step, we downloaded all models from the JWS database (date of download: June 17, 2019). As these models were comparably small, we complemented this collection by a set of larger models from the BioModels database (date of download: June 24, 2019) and a particularly large SBML model of cancer signalling [Frohlich et al., 2018], available at https://github.com/ICB-DCM/CS_Signalling_ERBB_RAS_AKT (commit 365e0be, date of download: June 24, 2019). These larger models allowed us to also assess the scaling behavior of computation times with respect to the size of the ODE models.

After importing these models with AMICI (version 0.10.7, commit d78010f), we assessed the correctness of the simulated state trajectories based on reference trajectories. For the models from JWS, reference trajectories were generated with the JWS built-in simulation routine, comprising 100 simulation time points along the trajectory. For the models from BioModels, we generated reference trajectories using the COPASI toolbox [Hoops et al., 2006]. For each model, the error tolerances were set to the strictest value which still allowed a successful integration of the ODE system. As the COPASI simulator is probably one of the the most widely used ODE solver implementation in the field of systems biology, we considered this the most reliable solution. Additionally, state trajectories were also compared by visual inspection.

All models were then simulated with AMICI and default settings - the Newton-type non-linear solver, the linear solver KLU, the BDF algorithm, and a maximum number of ODE solver steps of 10^4^ - for different error tolerances. We also tested whether a higher maximum number of ODE solver steps affected the result, but did not find a substantial impact. A model was added to the benchmark collection if for each time point of each simulated state trajectory, the absolute or relative error (i.e., the mismatch between the reference trajectory and the simulation) was below a predefined acceptance threshold. We investigated the number of accepted models in dependence of the acceptance threshold for various absolute and relative error tolerances (Supplementary Figure S1). For too strict acceptance thresholds, all models were rejected, but when simulating with high accuracy, gradually increasing the threshold led to a plateau of accepted models for thresholds between 10^−5^ and 10^−3^. When increasing the acceptance threshold further, the number of accepted models grew until eventually all models were accepted. We interpreted the plateau of models at threshold values around 10^−4^ as those models which could be simulated correctly with sufficient accuracy and hence fixed the final acceptance threshold to a value of 10^−4^.

This procedure resulted in accepting a total of 185 SBML models from JWS, out of 417 which could be successfully imported with AMICI. After grouping SBML models which belonged to the same biological model (i.e., which were grouped in one SED-ML [Waltemath et al., 2011] file on JWS), we were left with 141 independent models from JWS. Among the initially imported 30 models from the BioModels database, 25 were accepted. Together with the additional large-scale cancer signalling model, this left us with a total of 167 ODE models for our studies.

## Acknowledgements

We want to thank Jacky Snoep for his help and advice when working with the JWS online database of SED-ML and SBML models.

## Author information

### Contributions

P.S. wrote the implementation and performed the study under the direct supervision of Y.S. and L.S.. P.S., Y.S., L.S. and P.L.S. worked out the technical details. J.H. and P.L.S. conceived the study. All authors discussed the results and conclusions and jointly wrote and approved the final manuscript.

### Corresponding author

Correspondence to Jan Hasenauer.

