## Supplementary Information for "Benchmarking of numerical integration methods for ODE models of biological systems"

August 2020

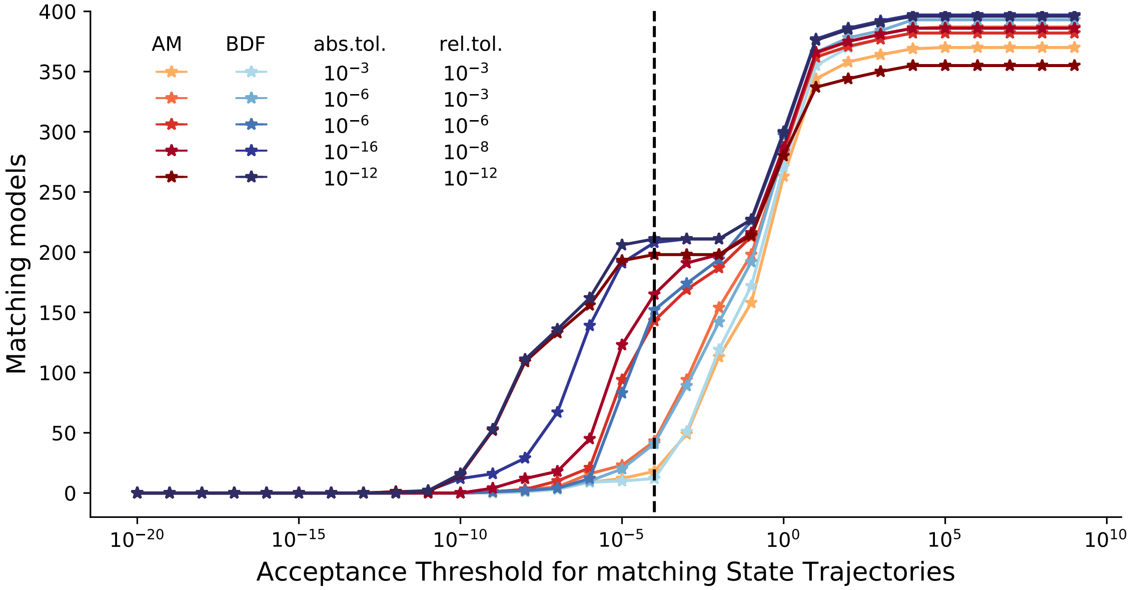


Figure S1: **Model Collection.** Number of accepted models with matching state trajectories for different acceptance thresholds. All models were simulated with AMICI using integration algorithms Backward Differentiation Formula (BDF) or Adams-Moulton (AM), with Newton-type non-linear solver, linear solver KLU, and six combinations of relative and absolute tolerance as depicted. Model simulations were compared to reference trajectories either from JWS or created with COPASI using the strictest tolerances which still allowed a model successful simulation.

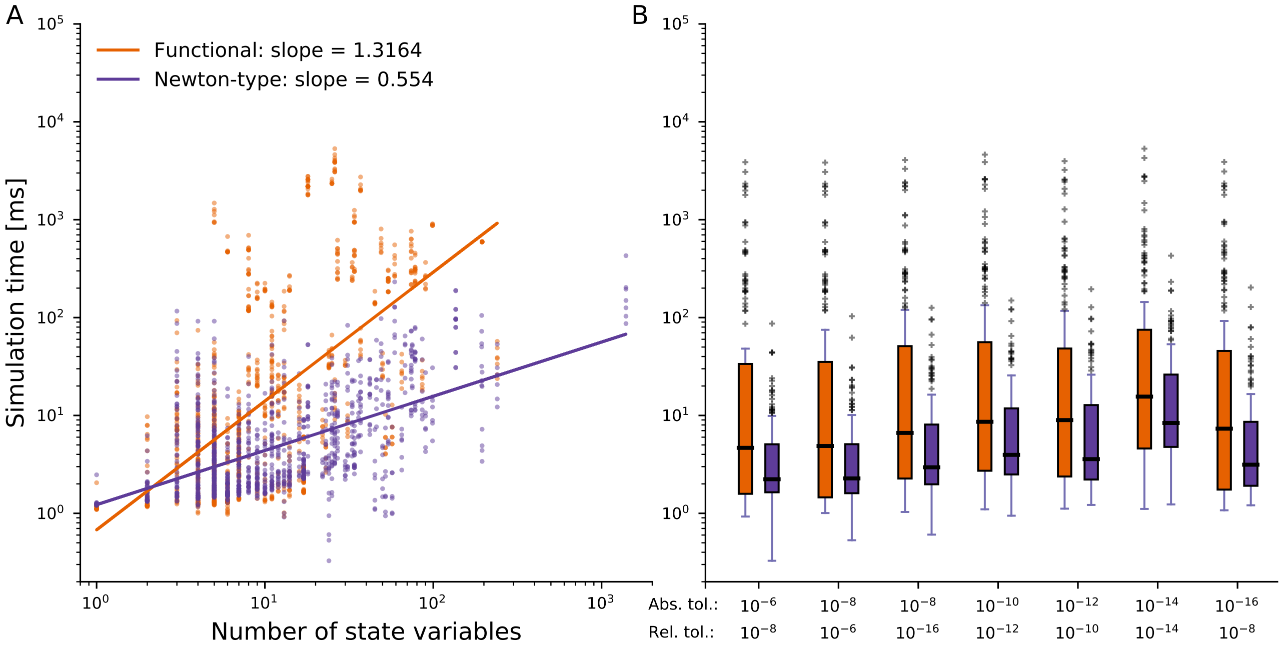


Figure S2: **Non-linear Solver.** Comparison between functional and Newton-type non-linear solver. **A** Scaling behavior of simulation times. Each point represents a model with a solver setting using the AM integration algorithm, the linear solver KLU, one out of two non-linear solvers and one out of seven tolerance combinations. Colors are used to visualize the different linear solvers. Colors are used to separate the two non-linear solvers The accompanying linear regressions depict the overall scaling behavior of computation time with respect to the number of state variables. The accompanying linear regressions depict the overall scaling behavior of computation time with respect to the number of state variables. **B** Computation time per tolerance combination. Visualization of the simulation time distribution using a setting of one non-linear solver in combination with seven tolerance combinations, the BDF integration algorithm and the KLU linear solver.

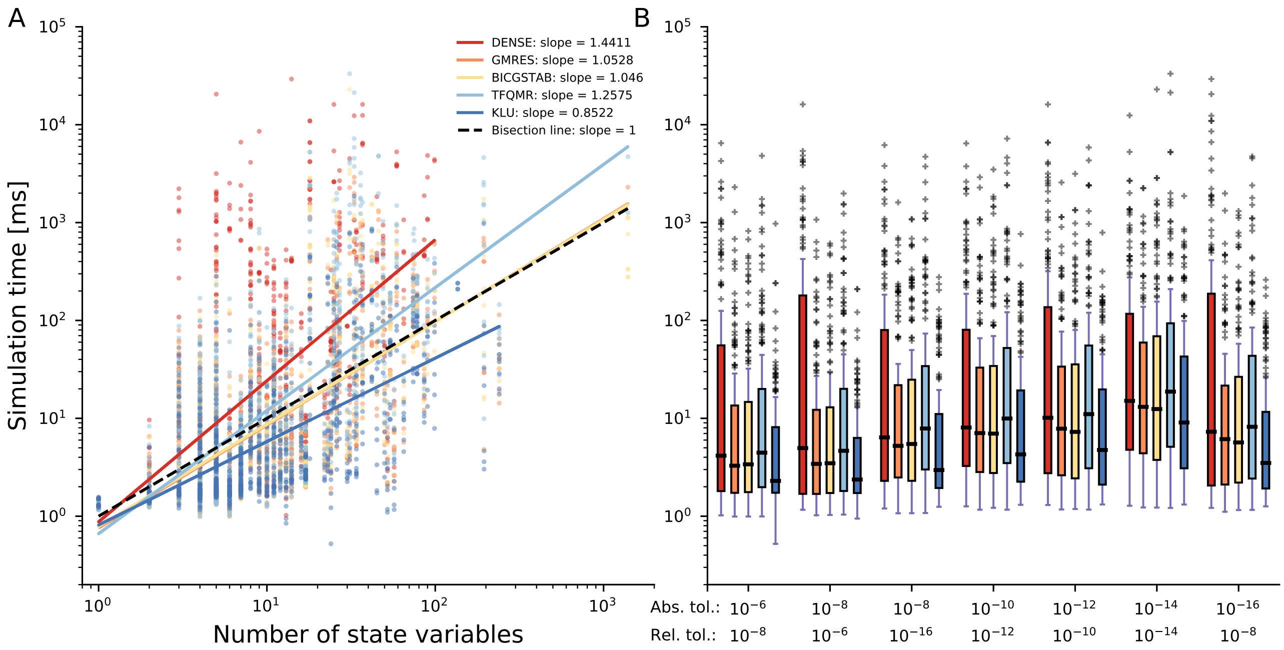


Figure S3: **Linear Solver using AM.** Comparison of the five linear solvers. **A** Scaling behavior of simulation times. Each point represents a model with a solver setting using one out of five linear solvers, AM integration algorithm, Newton-type non-linear solver and one out of seven tolerance combinations. Colors are used to visualize the different linear solvers. The accompanying linear regressions depict the overall scaling behavior of computation time with respect to the number of state variables. **B** Computation time per tolerance combination. Visualization of the simulation time distribution using a setting of one linear solver in combination with seven tolerance combinations, the AM integration algorithm and the Newton-type non-linear solver.

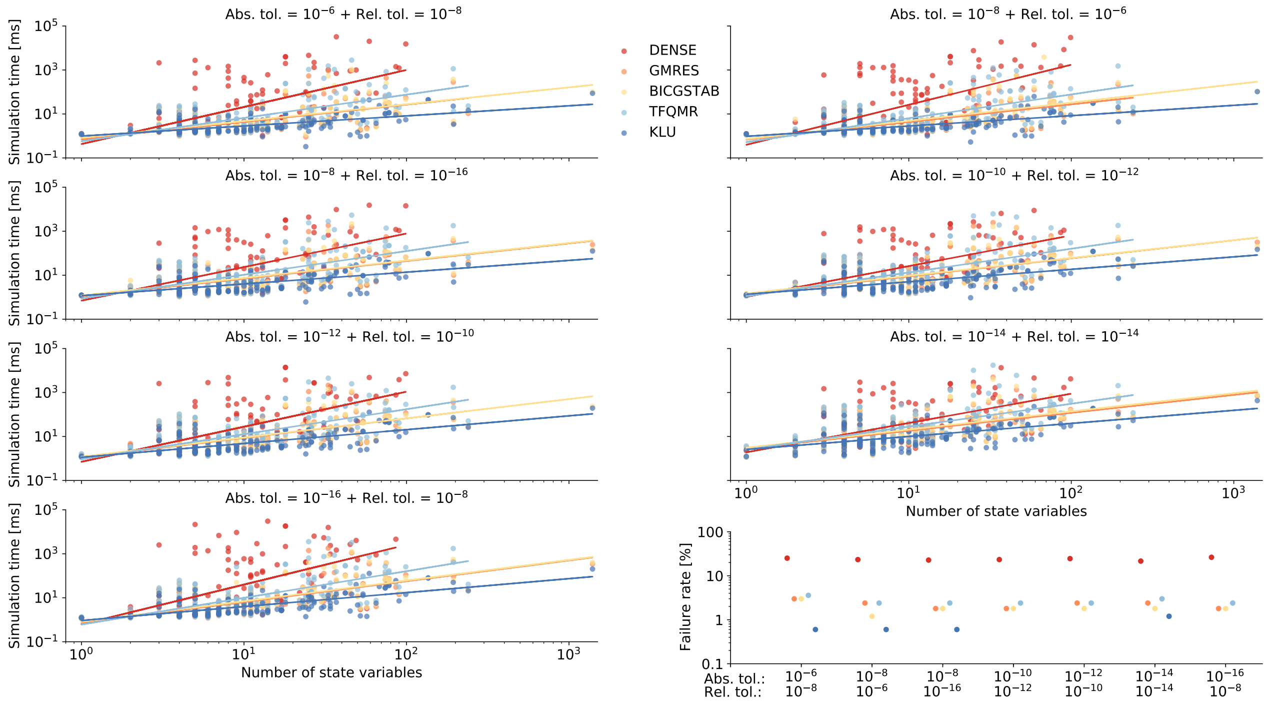


Figure S4: **Single linear solvers.** Visualization of simulation times. Each color represents a linear solver setting with Newton-type non-linear solver and BDF integration algorithm and one of seven tolerance combinations. The accompanying linear regressions display the scaling behavior with respect to the number of state variables. The last plot shows the failure rates of the linear solvers for all seven tolerance combinations.

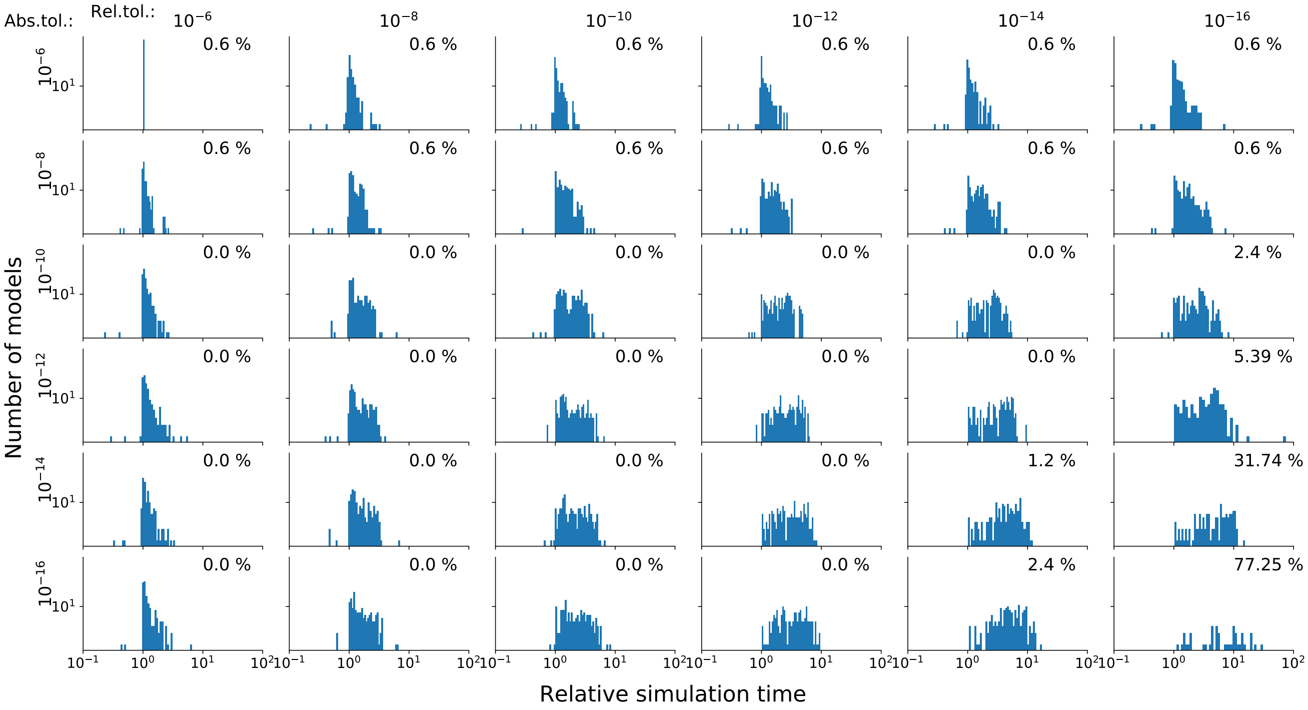


Figure S5: **Error tolerances.** Comparison of 36 tolerance combinations composed of absolute and relative error tolerances. The simulation time of each model was normalised by the simulation time using the laxest combination. The resulting simulation time ratios are depicted as histograms. The failure rate is displayed for each simulation setting in the top right corner. All models were simulated by using the Newton-type non-linear solver, the linear solver KLU and the integration algorithm BDF.

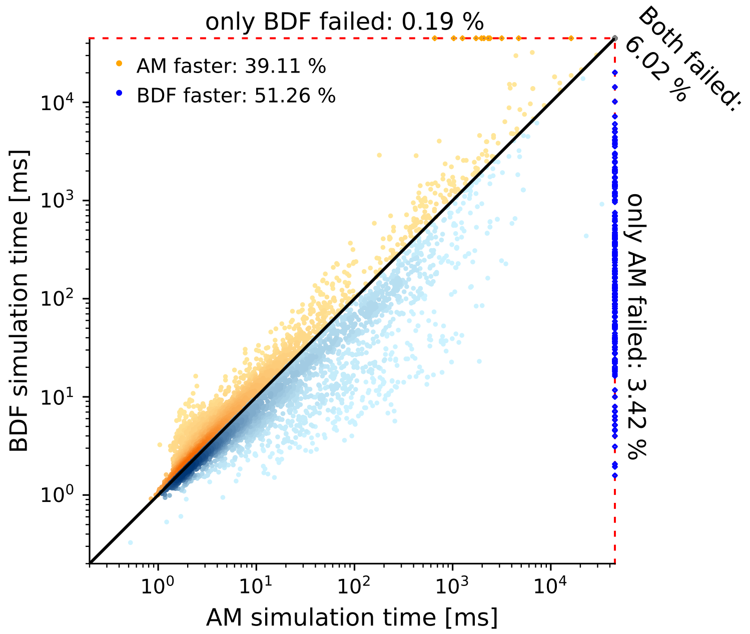


Figure S6: **AM vs BDF with all linear solvers.** Comparison between AM and BDF integration algorithm in terms of success rate and computation time. Each scatter point represents the computation time of a model by AM (x-axis) as well as BDF (y-axis) in combination with the Newton-type non-linear solver, one out of all linear solvers and the seven original tolerance combinations. The scatter points on the red dashed lines on top and on the right show failed simulations for one or both integration algorithms. Darker colors indicate a higher scatter point density.

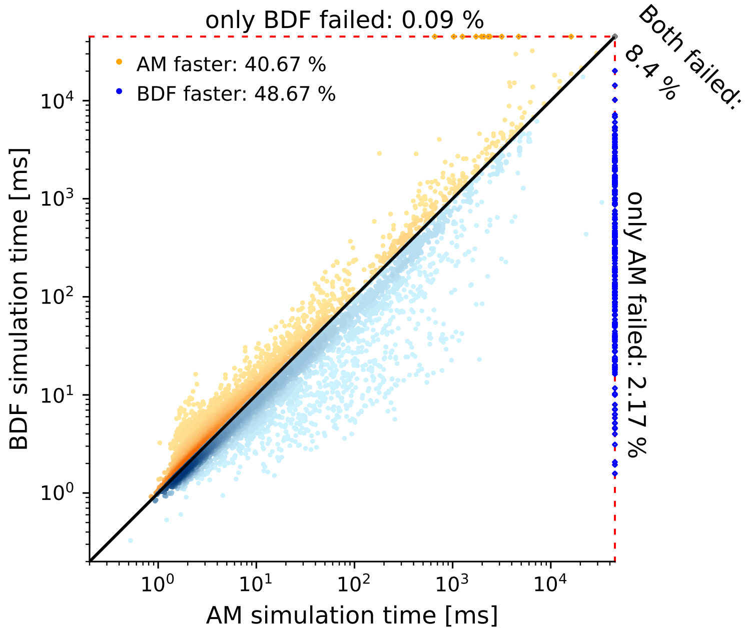


Figure S7: **AM vs BDF with all settings.** Comparison between AM and BDF integration algorithm in terms of success rate and computation time. Each scatter point represents the computation time of a model by AM (x-axis) as well as BDF (y-axis) in combination with one out of both non-linear solvers, all linear solvers and the seven original tolerance combinations. The scatter points on the red dashed lines on top and on the right show failed simulations for one or both integration algorithms. Darker colors indicate a higher scatter point density.
